# Control becomes habitual early on when learning a novel motor skill

**DOI:** 10.1101/2022.04.28.489941

**Authors:** Christopher S. Yang, Noah J. Cowan, Adrian M. Haith

## Abstract

When people perform the same task repeatedly, their behavior becomes habitual, or inflexible to changes in the goals or structure of a task. While habits have been hypothesized to be a key aspect of motor skill acquisition, there has been little empirical work investigating the relationship between skills and habits. To better understand this relationship, we examined whether and when people’s behavior would become habitual as they learned a challenging new motor skill. After up to ten days of practice, we altered the structure of the task to assess whether participants would flexibly adjust their behavior or habitually persist in performing the task the way they originally learned. We found that participants’ behavior became habitual early in practice—after only two days—at which point they were still relatively unskilled. These data demonstrate that motor skills become habitual after relatively little training, but can nevertheless further improve with practice.

## Introduction

We have all experienced the frustration of having to overcome old habits when we need to alter the way we perform a task. In a recent striking example of this, YouTuber Destin Sandlin created a “backwards bicycle”, a bicycle where rotation of the handlebar in one direction causes the front tire to rotate in the opposite direction (i.e., opposite of a normal bicycle) [1]. Although it is easy to understand how the handlebar moves the tire and it is trivial to rotate the handlebar, people find it difficult to ride the backwards bicycle, seemingly because they habitually try to balance themselves using the same movements they would perform on a normal bicycle.

In neuroscience and psychology, habits are generally defined as behaviors which, through extensive repetition, have become inflexible to changes in the goals or structure of a task [2–4]. Habit formation has almost exclusively been studied in the context of *discrete* choices [5–10], such as deciding which button to press on a keypad, or whether or not to perform an action at all [7, 11]. In such cases, habits are conceptualized as stimulus–response associations that have become obligatory through repetition [12–15].

Perhaps surprisingly, habit formation has hardly been studied in the context of *continuous* motor skills like riding a bicycle. In this case, the analog of a stimulus–response association guiding behavior is a *controller*, a mapping from the instantaneous states of the environment and one’s body to outgoing motor commands. Under this framework, behavior can be considered to be habitual if one’s controller for a task becomes habitual. That is, if the mapping between states and motor commands becomes inflexible to change and one persists in using this controller even if it no longer successfully achieves control objectives. Although it is conceptually straightforward to extend the concept of a habit from discrete tasks to continuous movement control, it is by no means clear that habits form in the same way in both cases. A key tenet of the stimulus– response framework is that a particular stimulus and resulting response must be paired repeatedly for a habit to form, but in continuous control tasks there are a continuum of (i.e., infinitely many) possible states and actions and it is unclear whether one will ever repeat the same action in the same state often enough for a habit to form.

Nevertheless, habits have been hypothesized to be a key aspect of motor skill acquisition, enabling more rapid behavioral responses and freeing up cognitive resources [16–20]. However, the exact relationship between habits and skill remains unclear [21], and progress in our understanding of this relationship is hampered by a dearth of empirical evidence examining how quickly behavior becomes habitual when learning a new motor skill. To a limited extent, behavior which could be interpreted as habits has been studied in continuous control tasks such as reaching under mirror-reversed visual feedback [22, 23] or more real-world skills like javelin throwing [24], swimming [25], and weightlifting [26]. However, such work has examined the process of replacing already highly skilled movements (baseline or well-practiced behavior) with new movements rather than how habits form when people *initially* learn a skill.

We performed an experiment to examine the process of habit formation in the context of learning a new continuous motor skill. Participants learned to control an on-screen cursor using a non-intuitive bimanual mapping where vertical movements of the left hand were mapped to horizontal movements of the cursor while horizontal movements of the right hand were mapped to vertical movements of the cursor. Previous work suggests that people learn this mapping by building a new controller *de novo* [27]—in contrast to how people learn simpler perturbations like rotations of visual feedback by adapting an *existing* controller [28–31]. Three separate groups of participants learned to control the bimanual mapping over two, five, or ten days of practice. At the end of the final day of practice, we flipped the direction of the mapping between movement of the left hand and movement of the cursor (i.e., a mirror reversal) and assessed whether participants would habitually continue to control the cursor according to the originally practiced mapping, or whether they would be able to flexibly adjust their control to accommodate the new flipped mapping.

Previous studies of habit formation have suggested that the expression of habits may be masked by deliberative, goal-directed processes that might override habitual responses during the reaction time prior to movement, particularly if participants are allowed ample time to prepare their movements [32]. To account for habitual control potentially being masked in this way, we assessed habitual behavior using two different approaches with differing reaction time constraints. In the first task, participants made point-to-point reaches towards targets in random locations, and they were allowed unlimited time to deliberate about each reach. In the second task, participants tracked a target moving in an unpredictable, sum-of-sinusoids trajectory. Here, the amount of time participants had to prepare their movements varied with the frequency of the target’s movement, with movements at high frequencies requiring particularly rapid responses and therefore minimal scope for deliberation.

Using these tasks, we sought to examine whether or not habits would be observable under the flipped mapping and, if so, when in the course of learning they would emerge.

## Results

### Participants became more skilled in the point-to-point task with practice over multiple days

Three different groups of participants learned to control an on-screen cursor using a novel bimanual hand-to-cursor mapping over two (*n* = 13), five (*n* = 14), or ten (*n* = 5) days of practice (two different versions of the bimanual mapping were used for different subsets of participants to control for potential biasing effects of biomechanics; see “Tasks” section of Methods). Participants practiced this mapping by performing a series of 12 cm point-to-point reaches towards targets, following a random walk within a 20 *×* 20 cm workspace. They alternated this point-to-point task with a tracking task in a block-wise manner (Fig 1).

**Fig 1.**
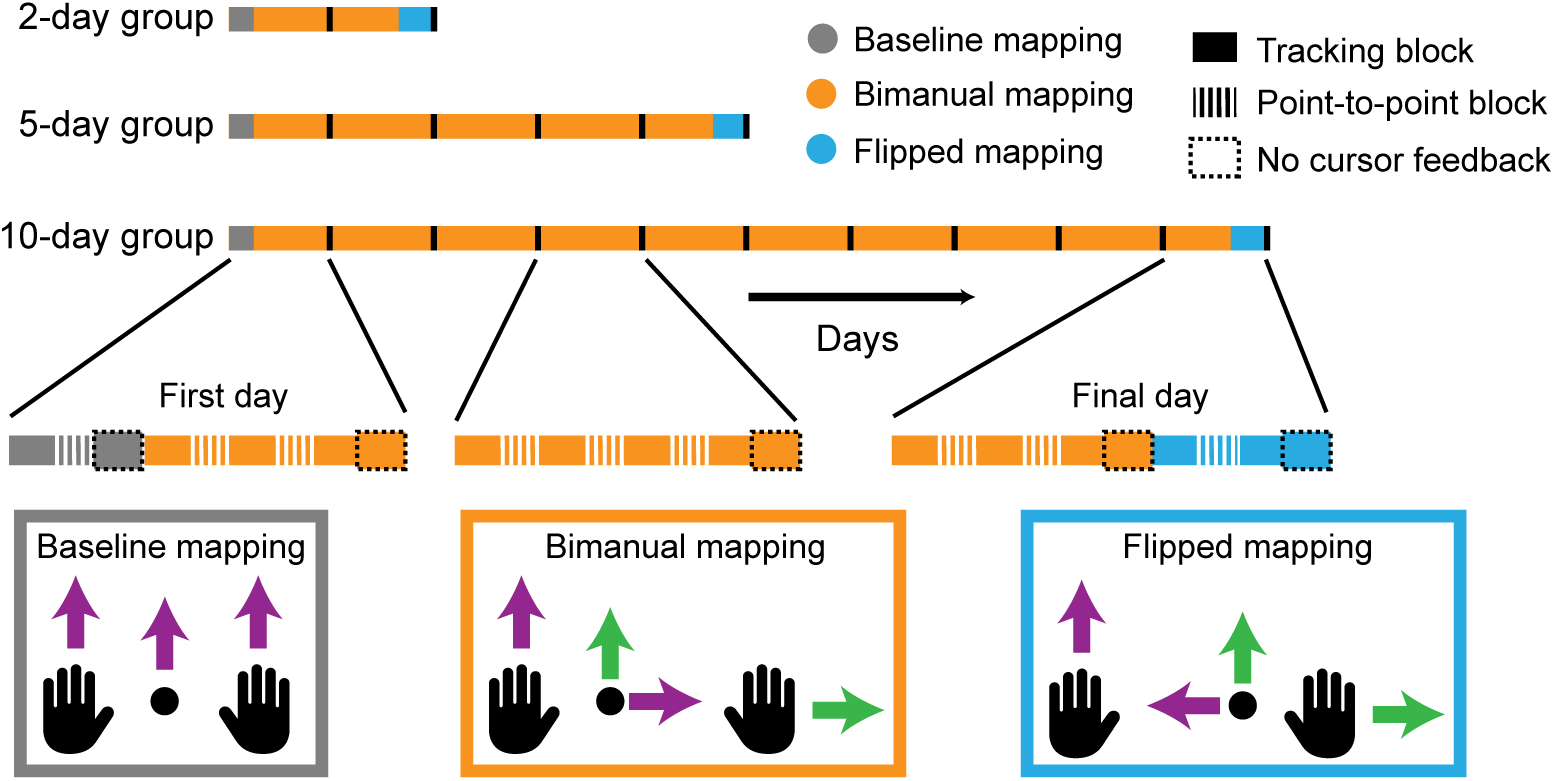
Tasks and experiment. Participants learned to control an on-screen cursor using a bimanual hand-to-cursor mapping (orange) over two (*n* = 13), five (*n* = 14), or ten (*n* = 5) days of practice. Half of the participants in each group practiced the depicted bimanual mapping while the other half practiced an alternate version where cursor movements were rotated 180^*°*^ relative to the depicted mapping (this was done to counterbalance any effects of biomechanics). On each day, participants performed blocks of point-to-point reaching (hashed rectangle; 1 block = 100 trials) and continuous tracking (1 block = 5 minutes) both with (solid rectangle) and without (solid rectangle with dashed outline) visual feedback of the cursor. Learning was compared relative to a baseline mapping where the cursor was placed at the average position of the two hands (gray). At the end of each group’s final training day, we flipped the left hand’s mapping to cursor movements (blue) and assessed whether participants would habitually continue to control the cursor under the bimanual mapping they originally learned.

Fig 2A shows representative raw cursor trajectories at baseline, early learning, and at late learning for each group in the point-to-point task. As previously found [27], participants initially experienced great difficulty in coordinating their two hands together to move the cursor straight towards each target. But they gradually improved their performance with practice, eventually moving between targets in a straight line, similar to their performance when using an easy mapping in which the cursor appeared exactly halfway between the left and right hand (“baseline”; Fig 1).

**Fig 2.**
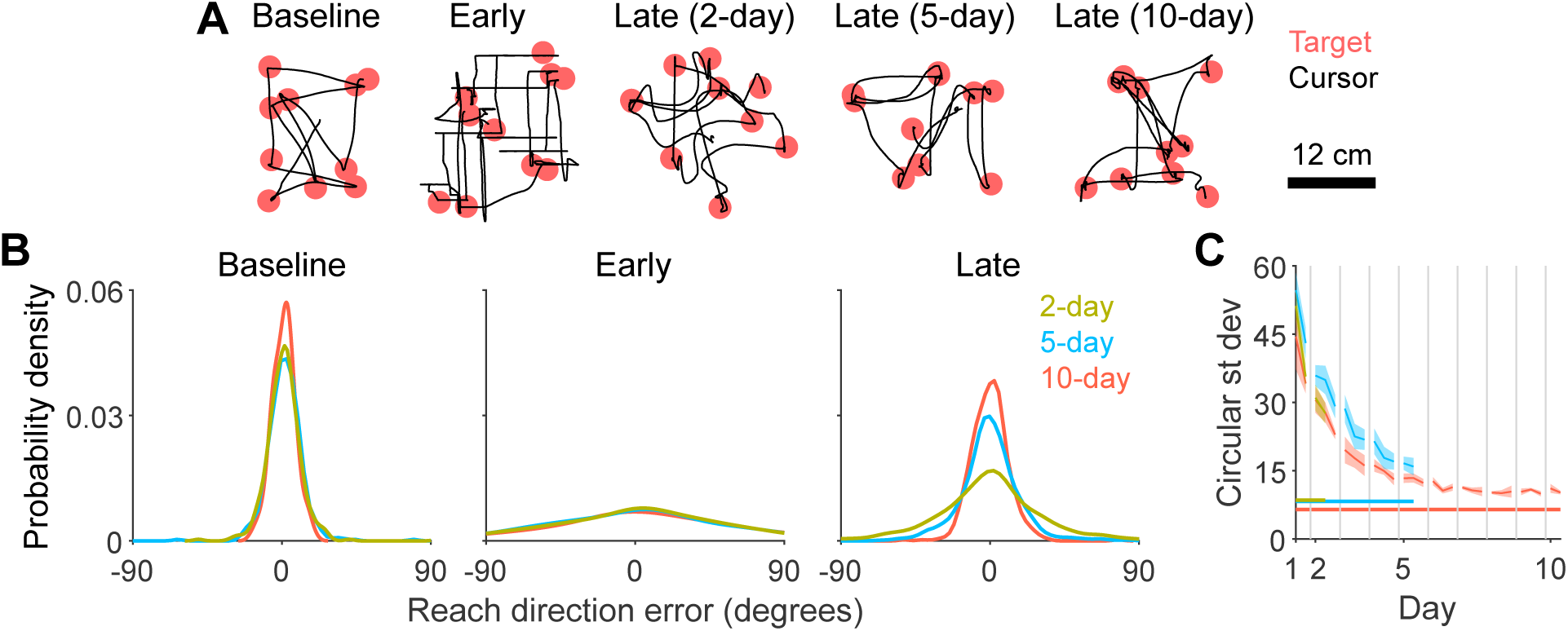
Performance in the point-to-point task under the bimanual mapping. A: Examples of raw cursor trajectories (black line) from baseline, early learning, and late learning (last block before flip block). Targets are displayed as red circles. Data from ten trials are shown for each block. B: Kernel-smoothed probability density of reach direction errors pooled over all subjects and trials for a given block. All blocks were the same as those shown in Fig 2A. C: Circular standard deviation of reach direction errors, computed by fitting a mixture model to the data in Fig 2B (see “Analysis of point-to-point task” section of Methods for more details). Each point corresponds to data from a single block and error bars indicate SEM across participants. Baseline standard deviations for each group are shown as horizontal lines. Days are demarcated by gray vertical lines.

We assessed each groups’ skill level by quantifying how precisely they aimed the cursor’s initial movement towards the target (Fig 2B-C; see Supplementary Fig 1 for other kinematic metrics throughout learning). Precision improved with practice, reaching a plateau after roughly 5 days that was close to baseline levels. Although there were small improvements in performance from day 5 to day 10 in the 10-day group, these improvements were not statistically significant (linear mixed effects model with post-hoc Tukey test [see Methods for details about statistical analyses]: *t* = *−*0.79, *p* = 0.9665). Thus, participants became more skilled in performing point-to-point reaches under the bimanual mapping, and the bulk of this improvement occurred over the first five days of practice.

### Performance in the tracking task also improved with practice over multiple days

Participants also performed a second task under the bimanual mapping where they tracked a target moving in a sum-of-sines trajectory (frequencies ranging from 0.1–1.85 Hz; Fig 3A). Unlike the point-to-point task where participants had an unlimited amount of time to plan their movements at the start of each trial, in the tracking task, the target moved quickly and pseudorandomly, limiting the amount of time participants had to plan their movements; any movements planned at one moment would become outdated within tens of milliseconds as the target would move to a new, unpredictable location. Additionally, the amount of time participants had to prepare their movements depended on the frequency at which the target moved. At low frequencies, the target oscillated slowly, providing people ample time to prepare their movements. But at high frequencies, the target oscillated quickly, forcing people to respond quickly (similar in effect to other approaches to limiting preparation time, such as forcing responses at particular time intervals [32]).

**Fig 3.**
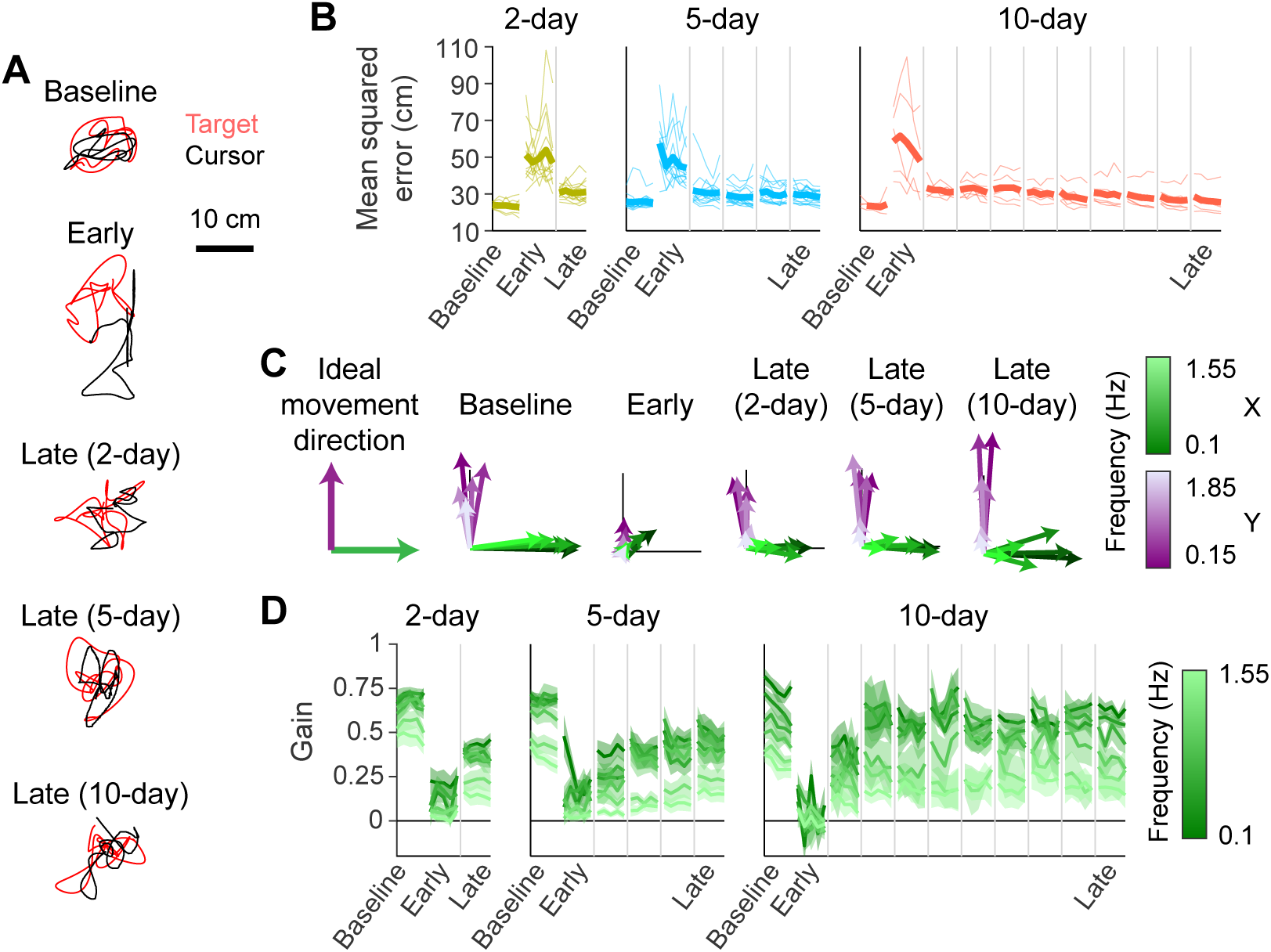
Performance in the tracking task under the bimanual mapping. A: Example cursor (black) and target (red) trajectories from single trials. B: Mean-squared error between cursor and target positions. Thin lines indicate individual participants while thick lines indicate group averages. Data collected from different days are separated by gray lines. Data from only one or two blocks are shown for each day for ease of visualization. C: Visualization of the cursor’s movement direction and gain (relative to the target) at frequencies of *x*- (green) and *y*-axis (purple) target movement. Each arrow depicts the direction and gain averaged across participants at a single frequency. Lower and higher frequencies are depicted as darker and lighter colors, respectively. Black lines are scale bars indicating a movement gain of 0.5. Visualization of ideal movement direction is shown on the left. D: Gain of horizontal cursor movements at frequencies of *x*-axis target movement (horizontal component of green arrows in Fig 3B). Error bars indicate SEM across participants.

With practice, participants learned to reduce the positional error between the target and cursor (Fig 3B). We examined participants’ tracking performance at different frequencies of movement using a system identification approach [33–37], which allowed us to separately examine the behavioral responses to target movements at different frequencies even though they occurred concurrently in the task. Specifically, we computed the gain and direction of cursor movement relative to target movement at each frequency, which can be interpreted analogously to the reach direction analysis for the point-to-point task in Fig 2B (see Methods for more details). Each arrow in Fig 3C shows the gain and direction of cursor movements in response to target movement at a particular frequency. Ideally, participants would track the target by moving their cursor in the same direction as the target. Thus, cursor responses to positive *x*-axis target movement (green) should be pointed rightwards while responses to positive *y*-axis target movement (purple) should be pointed upwards, which indeed was the case at baseline. By late learning, all groups exhibited movement gains and directions that approached that of baseline performance.

To statistically compare each group’s performance, we computed the gain of horizontal cursor movements at the frequencies of *x*-axis target movement (*x*-component of green arrows in Fig 3C). Gains improved from day 1 to day 2 in the 2-day group (Fig 3D; linear mixed effects model with post-hoc Tukey test; *p <* 0.05 for 4 out of 6 frequencies) and from day 2 to day 5 in the 5-day group (*p <* 0.05 for 3 out of 6 frequencies). However, gains did not significantly improve past day 5 in the 10-day group (*p >* 0.05 for all frequencies). These data demonstrate that, with practice, participants became able to successfully move their hands in the appropriate direction to track the target and, as in the point-to-point task, the bulk of this improvement occurred in the first 5 days.

### Participants’ behavior in the point-to-point task was habitual after only two days of practice

Having examined participants’ skill, we next asked whether and when their behavior became habitual. Might participants’ behavior become habitual around the same time their skill plateaued (i.e. by Day 5), early in learning (i.e. by Day 2), or only after significant repetition of the stably performed skill (i.e. by Day 10)? Or, lastly, might participants behavior never have become habitual? To determine this, at the end of each groups’ final day of practice, we had participants control the cursor under a new flipped mapping where the mapping between the left hand and the cursor movement was reversed relative to what they had originally practiced (“flip” block), effectively amounting to a left-right mirror reversal applied on top of the originally practiced bimanual mapping. Participants were explicitly informed about the reversal of their left hand’s mapping, and we tested whether participants would habitually continue to control the cursor under the originally learned bimanual mapping or successfully alter their behavior according to the flipped mapping.

First, we assessed whether participants exhibited habitual behavior in the point-to-point task. On a given trial, if participants could successfully control the cursor under the flipped mapping, then we would expect their cursor’s initial movement to be aimed towards the true target (i.e., goal-directed). But if participants habitually controlled the cursor according to the original bimanual mapping, then we would expect their cursor’s initial movement to be aimed towards a virtual target reflected directly across a vertical mirroring axis. We found that participants in all three groups exhibited both goal-directed and habitual behavior during the flip block (Fig 4A) on different trials. We visualized how often participants’ movements were aimed towards the virtual mirrored target as a heat map plotting the cursor’s initial movement directions as a function of the target’s direction (Fig 4B). If participants reached towards the virtual mirrored target, their initial cursor directions would lie along the *y* = *−x* line. Although none of the groups exhibited initial cursor directions along this line at late learning, all groups did exhibit such behavior during the flip block.

**Fig 4.**
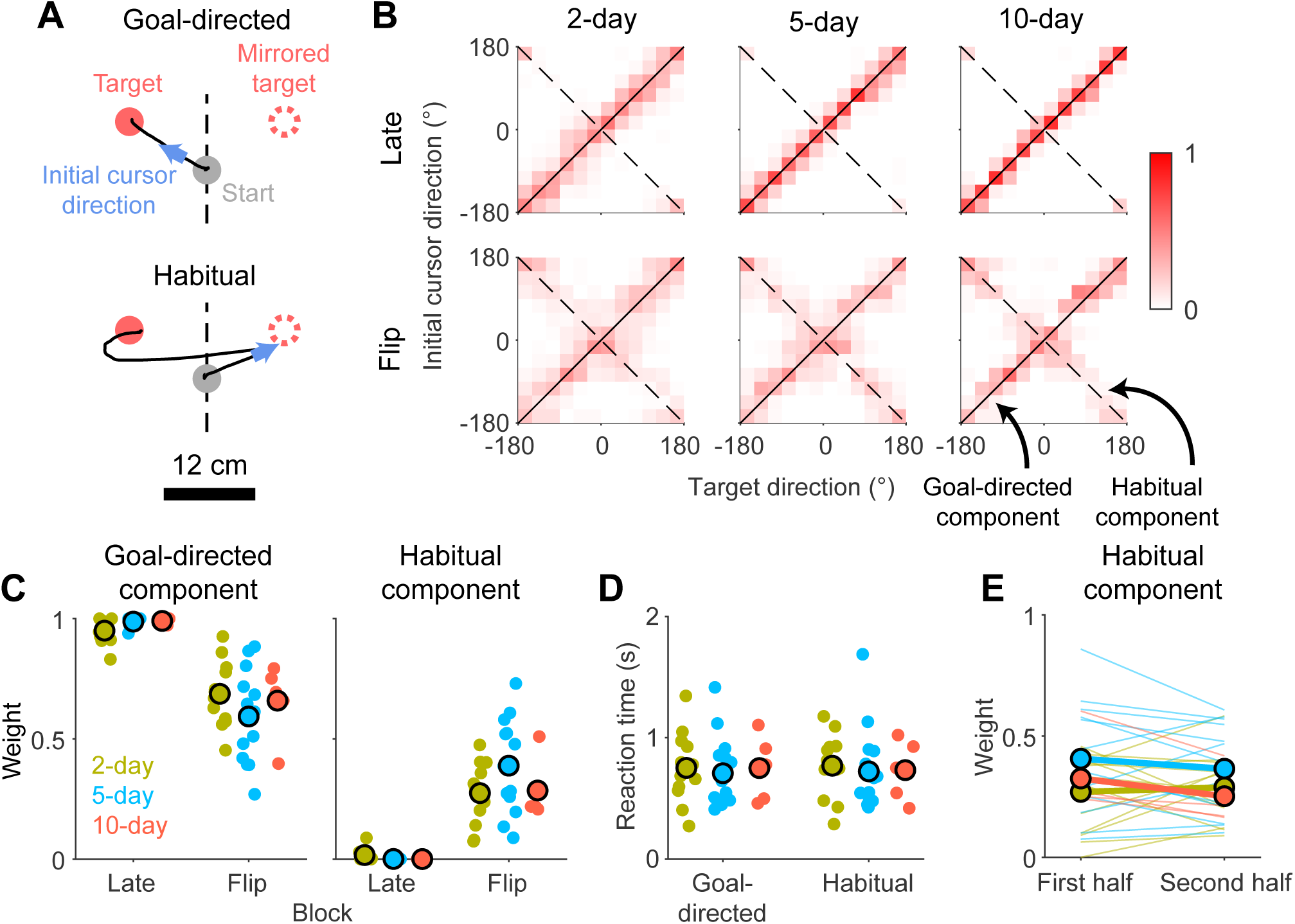
Analysis of habit in the point-to-point task. A: Cursor trajectories (black line) from single trials in the flip block. The trajectories show trials where the cursor’s movement was initially aimed straight towards the target (upper) or aimed towards a virtual target (lower) mirrored across the vertical axis (dashed line). B: Heat map of cursor’s initial movement direction as a function of target directions. Data were pooled from all subjects and grouped into bins of 30^*°*^ on both axes. We defined 0^*°*^ as the positive *y*-axis (i.e., the mirroring axis). Within each target direction bin, we computed the fraction of trials which fell in a particular reach direction bin, plotting this fraction as color intensity in the heat map. To measure the proportion of trials where participants exhibited habitual behavior, we fit a mixture model composed of two weighted von Mises distribution centered on either the *y* = *x* (goal-directed) or *y* = *−x* line (habitual behavior). C: Fitted weights for the goal-directed (left) and habitual (right) components of the mixture model depicted in Fig 4B. Fits for individual participants shown as small circles and group means shown as large circles. D: Reaction time for reaches towards the actual target (goal-directed) versus a virtual mirrored target (habitual). E: Weight of the habitual component of the mixture model when fit to data from either the first or second half of the flip block. Thin lines are individual participants and thick lines are group means.

We estimated the proportion of trials where participants initially reached towards the mirrored target by fitting a mixture model to the reach direction data (see “Analysis of point-to-point task” section in Methods for more details; see Supplementary Fig 2 for a model comparison and model recovery analysis), using this as a metric for how strongly participants exhibited habitual behavior. We found that the proportion of habitual movements was significantly higher in the flip block compared to late learning for all three groups (Fig 4C; linear mixed effects model with post-hoc Tukey test; 2-day: *t* = *−*5.89, *p <* 0.0001; 5-day: *t* = *−*9.18, *p <* 0.0001; 10-day: *t* = *−*4.61, *p* = 0.0010). These data demonstrate that all groups exhibited habitual behavior in the point-to-point task.

Perhaps surprisingly, the proportion of reaches towards the mirrored target was not significantly different between groups (Fig 4C; linear mixed effects model with post-hoc Tukey test; 2-day vs. 5-day: *t* = 2.08, *p* = 0.3098; 2-day vs. 10-day: *t* = *−*0.46, *p* = 0.9973; 5-day vs. 10-day: *t* = 1.08, *p* = 0.8887). In other words, despite the fact that the 10-day group practiced using the original bimanual mapping for five times as long as the 2-day group, they did not exhibit more strongly habitual behavior in the point-to-point task. Moreover, the reaction times for goal-directed reaches were not significantly different from habitual reaches (Fig 4D; linear mixed effects model; no main effect of group [*F* (2, 29) = 0.09, *p* = 0.9168], reach [*F* (1, 29) = 0.03, *p* = 0.8705], or interaction [*F* (2, 29) = 0.09, *p* = 0.9167]), suggesting that the lack of differences across groups in Fig 4C could not be explained by differences in the amount of time participants had to plan their movements.

In the flip block, we noted that participants occasionally adopted a strategy of initially moving the cursor vertically (the axis along which the mapping had not changed) before initiating the horizontal component of their movement. Therefore, as an alternative assay for habitual behavior, we computed the proportion of trials where the horizontal component of the cursor’s movement was initially directed away from the target. This analysis yielded qualitatively similar results, with behavior in all three groups being habitual, and no significant differences in the strength of habits across groups (Supplementary Fig 3; linear mixed effects model with post-hoc Tukey test; 2-day vs. 5-day: *t* = *−*0.79, *p* = 0.9685; 2-day vs. 10-day: *t* = 0.30, *p* = 0.9996; 5-day vs. 10-day: *t* = *−*0.28, *p* = 0.9998). Collectively, these results suggest that in the point-to-point task, participants exhibited equally strong habitual behavior regardless of whether they had practiced using the bimanual mapping for two, five, or ten days of practice.

### Behavior in the tracking task also became habitual after only two days of practice

We next examined participants’ behavior in the tracking task to see whether they would exhibit similar habitual behavior under the flipped mapping, or whether habit effects might even be exacerbated given the imperative to generate movements rapidly while tracking the target. We compared the direction of participants’ responses (i.e. cursor movement) to movements of the target between late learning and the flip block (Fig 5A). During the flip block, if participants habitually behaved according to the original bimanual mapping, then their hand movement in response to horizontal target movement would be similar to late learning, and the horizontal movement of the cursor would therefore be directed opposite to the movement of the target. We expected that the extent of this effect might vary according to the frequency of target motion, with high frequencies being more likely to appear habitual owing to the need to respond more rapidly. We normalized the cursor’s horizontal movement gain from the flip block by the gain from late learning such that normalized gains of -1 would indicate habitual behavior (Supplementary Fig 4). (In subsequent analyses, we removed data from one outlier participant in the 10-day group—bottom left participant in Supplementary Fig 4—who exhibited erratic behavior in the flip block, with negative gains that were greater in amplitude than at late learning.) In the first flip block, all groups on average exhibited negative gains at two or more frequencies (Fig 5B; one-sample t-test with Holm-Bonferroni correction at *α* = 0.05; 2-day: 2 of 6 frequencies; 5-day: 3 of 6 frequencies; 10-day: 2 of 6 frequencies), particularly at higher frequencies, as we expected. However, we did not find any evidence that groups with more practice exhibited significantly more negative gains than groups with less practice (linear mixed effects model with post-hoc Tukey test: *p >* 0.05 for all comparisons of gains within frequencies).

**Fig 5.**
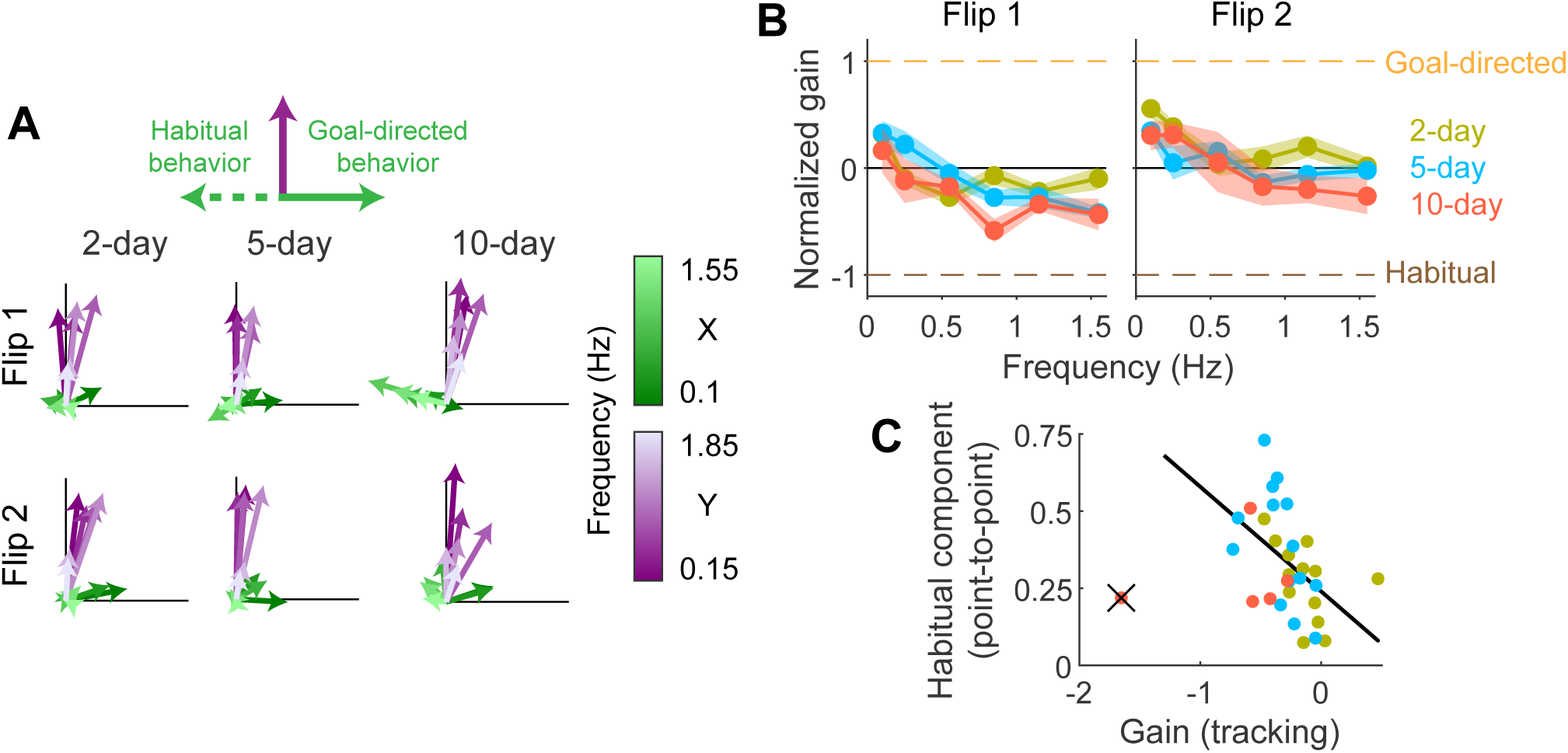
Analysis of habit in the tracking task. A: Visualization of the cursor’s movement direction and gain (relative to the target) while using the flipped mapping, similar to Fig 3B. Each arrow depicts the average across participants at *x*- (green) and *y*-axis (purple) frequencies. Lower and higher frequencies are depicted as darker and lighter colors, respectively. Black lines are scale bars indicating a movement gain of 0.5. Flip 1 and Flip 2 are the first and second tracking blocks under the flipped mapping, respectively. At the top, we depict the direction the green arrows should point if participants exhibit goal-directed (right) or habitual (left) behavior. B: Gain of horizontal cursor movements under the flipped mapping normalized to the gain under the original bimanual mapping at late learning. If participants responded to target movement by moving their cursor in the same direction under both the original and flipped bimanual mappings, then the gain would be positive, approaching 1 if the movement gains were the same (goal-directed; yellow dashed line). But if they moved their cursor in opposite directions under the two mapping, then the gain would be negative, approaching -1 (habitual; brown dashed line). Error bars indicate SEM across participants. C: Linear regression between average gains of the highest 3 frequencies from the first flipped tracking block and the proportion of habitual reaches from the flipped point-to-point block in Fig 4C. Data from one outlier subject in the 10-day group (crossed out in black) was not used for fitting.

The above analyses considers differences in behavior across groups. However, in previous work, we have found that habitual behavior can vary greatly across individuals [32]. We therefore also examined whether or not behavior was habitual at an individual-participant level. We calculated the proportion of participants in each group who exhibited significantly negative gains during the flip block. In all three groups, we found a mixture of habitual and non-habitual participants (Supplementary Fig 4; one-sample t-test with Holm-Bonferroni correction at *α* = 0.05; 2-day: 5 of 13 participants; 5-day: 7 of 14 participants; 10-day: 3 of 4 participants). Although the proportion of participants who were habitual did appear to increase with practice, it is difficult to conclude whether or not this trend was meaningful, particularly given the small sample size of the 10-day group.

### The strength of habitual behavior was correlated between the point-to-point and tracking tasks

Might there be any relationship between the habitual behavior we observed in the point-to-point and tracking tasks? To examine this, we compared how strongly participants exhibited habitual behavior between the two tasks. First, we averaged the gains of each participant’s tracking behavior at the highest three frequencies, given that we expected habitual behavior to be strongest at these frequencies [37]. We then correlated each subject’s average gain with the proportion of habitual reaches they made in the point-to-point task, as in Fig 4C. Indeed, we found a correlation between tasks (Fig 5C, slope = *−*0.34, Pearson’s *r* = 0.49, *p* = 0.0052), suggesting that the tasks may have indeed assessed the same underlying habit.

### All groups’ habits were similarly resistant to extinction

An alternative way in which behavior may become habitual is by becoming more persistent, i.e., resistant to extinction. In other words, as one’s skill increases, the habits one forms may persist for longer. To assess whether habitual behavior would became more persistent with more training, we examined participants’ performance in a second tracking block under the flipped mapping, after having practiced the flipped mapping in a point-to-point block Fig 1). At the group level, participants no longer exhibited significantly negative gains at any frequency (Fig 5B; one-sample t-test with Holm-Bonferroni correction at *α* = 0.05), suggesting that the habits had been largely extinguished in all groups. However, at the level of individual participants, all 3 of participants in the 10-day group who exhibited negative gains in the first tracking block still did so in the second tracking block. Meanwhile, only 1 of 5 of participants in the 2-day group and 2 of 7 of participants in the 5-day group still exhibited significantly negative gains. While these data suggest that habitual behavior may have been more persistent in the 10-day group, again, they must be interpreted with caution given the small sample size of this group.

We attempted a similar analysis of the persistence of participants’ habits by fitting the mixture model from Fig 4D to the first and last 50 reaches in the block instead of all 100 reaches. However, we found no evidence for any difference in the strength of habitual behavior between the first and last half of the block for all groups (Fig 4E; linear mixed effects model with post-hoc Tukey test; 2-day: *t* = *−*0.49, *p* = 0.9963; 5-day: *t* = 1.11, *p* = 0.8747; 10-day: *t* = 1.17, *p* = 0.8493), suggesting that this aspect of habitual behavior did not become extinguished over this period.

## Discussion

In the present study, we examined the time course over which habitual behavior emerged as participants learned a new continuous motor skill—controlling a cursor under a novel bimanual hand-to-cursor mapping. Participants became more skilled in using this mapping by practicing with a combination of point-to-point reaches and continuous tracking, and their skill level plateaued after around five days of practice. After either two, five, or ten days, we flipped the left hand’s control of the cursor and tested whether participants would habitually continue to control the cursor according to the original mapping they had learned. We found that habitual behavior emerged after only two days of practice, which we observed in both the point-to-point and tracking tasks. We did not find compelling evidence, however, that habitual behavior became stronger with more practice.

While we mainly focused on assessing how *strong* habitual behavior became during learning, to a limited extent, we also assessed how *persistent* habits became, examining whether habits were extinguished through exposure to the flipped mapping. While this short period of practice seemed to be sufficient to extinguish the habit in the tracking task, we did not observe extinction in the point-to-point task. It is unclear why we observed a difference between the two tasks; although we expected the strength of habitual behavior to be affected by the amount of preparation time participants were afforded in each task, we did not expect habit persistence to be affected. However, we emphasize that we did not find strong evidence to suggest that the persistence of habitual behavior depended on how long participants practiced the bimanual mapping.

It is important to note that when we analyzed individual participants’ behavior, we found that a greater proportion of participants exhibited habitual behavior with more practice. Therefore, taking a conservative interpretation, one could say that our results are inconclusive as to whether behavior became more habitual with more practice. However, the inferences we could draw from the individual participant analysis were limited because they critically relied on data from our most practiced group which had a small sample size (*n* = 5), and if one were to only use data from the other two larger groups, there is little evidence to suggest more participants in the 5-day group (*n* = 14) exhibited habitual behavior than the 2-day group (*n* = 13). Furthermore, our main result that the emergence of skill and habit dissociated during learning would not be impacted by the findings of the individual participant analysis.

One additional aspect of our experiment that we did not discuss (since it was not directly relevant to our primary question of when control became habitual) was that at the end of each day, participants performed an additional tracking block without visual feedback of the cursor’s position. We used this block to examine the extent to which participants’ learning could be attributed to improvements in feedforward control. However, we found that for all groups, there was negligible improvement in mean-squared tracking error throughout learning and movement gains remained low (Supplementary Fig 5), indicating that participants were not capable of expressing their learned behavior without visual feedback of the cursor.

Collectively, our results suggest a dissociation between the emergence of skill and habit during motor learning: behavior can become habitual early in learning before one’s skill level has asymptoted, and likewise, behaviors which have already become habitual can still become more skillful through practice. Our findings parallel that of [32] who demonstrated that participants learning an arbitrary visuomotor association task exhibited improved speed-accuracy tradeoffs (i.e., improved skill) over twenty days of practice, even after behavior had become habitual after four days of practice. They further found that the habits could be explained as an all-or-none phenomenon (i.e., one either is or is not habitual), consistent with our observation that habitual behavior did not become stronger with more practice. The skill in [32] is quite rudimentary in that performance improvements amount only to speeding up action selection by tens of milliseconds, which might potentially have occurred via a specialized mechanism. In the present study, however, the improvements in skill after becoming habitual were more significant and unlikely to be explained by improvements in the processing speed alone. The similarity of the results between the two studies suggest a potential commonality in the process by which habits form when learning new skill in both discrete and continuous domains.

Many studies of habits have suggested that habits may not be an all-or-nothing phenomenon, but that some habits could be stronger or weaker that others. It remains unclear, however, exactly what it means for a behavior to become more strongly habitual. Habit strength could correspond to the strength of its immediate influence on behavior, e.g. how likely a habit is to be expressed, or it could correspond to how much the habitual behavior persists, or to some other related feature of behavior (e.g. stereotypy of movement. What metrics should we use to quantify these characteristics of habits? In our study, we quantified habit strength differently between the point-to-point (probability of expressing a habitual reach) and tracking tasks (gain of habitual movement), though we found that these measures were correlated. While this might suggest that these two assays measured a single underlying in different ways, we did also observe a dissociation in the persistence of behavior measured by these two different assays. Characterizing the ‘strength’ of a habit is further complicated by the fact that multiple component processes/computations may be involved in generating movement, and any one of these processes may become habitual. For instance, one’s ability to select what action to do (e.g., move the cursor to the right) may become habitual independently of one’s ability to execute that action (e.g., stereotyped kinematics of rightward movement; see [21] for a more in-depth discussion of this idea).

The present study provides new and important empirical evidence regarding how habits form and their relationship to skill. Our findings also have important implications for theoretical accounts of habits. A wide variety of theories have been proposed to explain the cognitive basis of habits, such as forming stimulus– response associations [12–15], caching expected future rewards [38–40], and caching computations/policies [19,41]. Central to these theories is the idea that habitual behavior is inflexible to change. Though behavioral inflexibility is (rightly) central to the definition of habits, our findings suggest that one may not need to break a habit in order to alter habitual behavior. To account for this, theoretical accounts of habits should allow scope for habitual behaviors to remain flexible to some degree. Learning rules which might accomplish this are incorporated into reinforcement learning-based frameworks, in which habitual behavior is often identified with model-free reinforcement learning which can still learn from experience, albeit slowly [38–40]. Our results underscore that habitual behavior is not set in stone, but can continue to evolve with experience—in this case, over multiple days of practice, unlike in model-free reinforcement learning where it can be observed in single trials [42].

How could a behavior be altered yet simultaneously remain habitual? If it is true that individual behaviors are generated via multiple intermediate computations that can independently become habitual, then it is possible for one computation to become fixed (i.e., habitual) while a different computation continues to improve with practice (i.e., more skillful). For instance, participants in our experiment may have habitually chosen to use a controller for the bimanual mapping, but their movement execution having chosen this controller could still become quicker and more precise. Alternatively, it is possible that computations which have become habitual simply maintain some small level of flexibility within which the behavior can be altered, but this learning may be slow and limited in scope. Understanding how exactly flexibility arises in habitual behavior should be a focus of future work.

To conclude, a behavior becoming habitual is often viewed as the final step in learning: learned behavior must be repeated in order to render it habitual, at which point it becomes a persistent and dependable component of skilled performance. While this idea may seem intuitively true, it has not previously been empirically tested. Our results challenge this view, suggesting that behaviors become habitual early in learning but maintain some flexibility to change with experience. What do our results ultimately suggest about the role that habits play when we learn new motor skills? During learning, one may encounter situations where they must alter their behavior to improve task performance, and often these behaviors will have already become habitual through practice. Overcoming habits will likely be frustrating when one must substantially alter their behavior [43]. But if only slight alterations are needed, then one may be able to fine tune their habit without having to break it.

## Methods

### Participants

A total of 32 right-handed participants were recruited for this study (23.0 *±* 4.3 [mean *±* standard deviation]; 13 male, 19 female), 13 for the 2-day group, 14 for the 5-day group, and 5 for the 10-day group (recruitment for the 10-day group was cut short due to the COVID-19 pandemic). All participants reported no prior history of neurological disorders. All methods were approved by the Johns Hopkins School of Medicine Institutional Review Board and were carried out in accordance with relevant guidelines and regulations. Informed consent was obtained from all participants in the study.

### Experimental setup

Participants were seated in front of a table with both of their hands supported on the table by frictionless air sleds. The position of participants’ hands were monitored at 130 Hz using a Flock of Birds magnetic tracker (Ascension Technology, VT, USA) placed near each hands’ index finger. Participants viewed stimuli on a horizontal mirror which reflected an LCD monitor (60 Hz), and the mirror obscured vision of both hands.

### Tasks

Participants learned to maneuver an on-screen cursor (circle of radius 2.5 mm) using one of two versions of a bimanual hand-to-cursor mapping. Half of the participants learned one version where up-down movements of the left hand produced right-left movements of the cursor while right-left movements of the right hand produced up-down movements of the cursor. The other half of participants learned a different version where the mapping from hand to cursor movements were 180^*°*^ rotated relative to the previous version. We counterbalanced these two versions of the mapping to ameliorate any effects biomechanics may have had on our results.

Three different groups practiced the bimanual mapping over either two, five, or ten days of training. Training consisted of participants making point-to-point reaches towards randomly placed targets (circles of radius of 10 mm) within a 20 *×* 20 cm workspace. Participants were instructed to reach towards each target as quickly and accurately as possible, with each trial consisting of one target location. Once the cursor was stationary (speed *<* 0.065m*/*s) within the target for 1 second, the target appeared in a random direction 12 cm away. To encourage participants to move quickly, we provided feedback to the participants indicating whether their peak velocity exceeded 0.39 m/s on that trial. If this threshold was exceeded, the target turned yellow and a pleasant tone was played once the cursor reached the target, and if the threshold was not exceeded, then the target did not turn yellow and no tone was played. The baseline point-to-point block consisted of 30 trials while all other point-to-point blocks consisted of 100 trials.

In between blocks of point-to-point reaching, participants performed a manual tracking task. In this task, a target moved continuously on the screen in a sum-of-sinusoids trajectory. The trajectory was composed of twelve sinusoids, six each in the *x*- and *y*-axes, parameterized by amplitude, 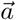, frequency, 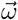, and phase, 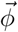, vectors. The target’s position along a single axis, *r*, was computed as:

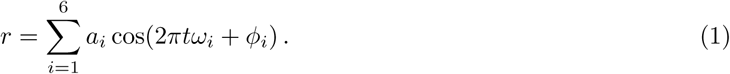

For the *x*-axis, 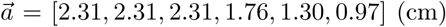 and 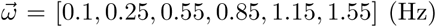. For the *y*-axis, 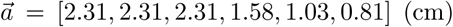 and 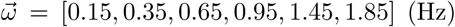. The values of 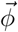 were randomized between [*−π, π*). Each trial lasted 66 seconds, and during the first 5 seconds of each trial, the target’s amplitude ramped up linearly from 0 to its full value. Each block consisted of 5 trials. Periodically, participants performed a tracking block without visual feedback of their cursor.

To assess the extent to which participants control of the bimanual mapping had become habitual, at the end of each groups’ final day of training, we flipped the left hand’s mapping to cursor movement (flip block): up-down movements of the left hand now resulted in left-right movements of the cursor instead of right-left (in the case of the 180^*°*^ rotated bimanual mapping, right-left movements of the cursor became left-right). We required three different groups of participants for this experiment because we assessed habitual behavior at different time points during learning, and after a participant has used the flipped mapping once, any future learning of the original bimanual mapping would be contaminated. The order of all blocks during the experiment is depicted in Fig 1.

## Data analysis

### Software

Data analyses were performed in MATLAB R2018b (The Mathworks, Natick, MA, USA) and R version 4.0.2 (RStudio, Inc., Boston, MA, USA; [44]) using the Matrix [45], lme4 [46], lmerTest [47], and emmeans packages [48]. Figures were generated using Adobe Illustrator (Adobe Inc., San Jose, CA, USA).

### Analysis of point-to-point task

The cursor’s position in each trial was smoothed using a third-order Savitzky-Golay filter. Path length was defined as the total distance that the cursor traveled in a single trial. Movement time was defined as the time between movement initiation (when the cursor left the start target) and termination (when the cursor was in the end target with speed *<* 0.065m*/*s). Reaction time was defined as the time between when the target appeared and the cursor’s tangential velocity exceeded 0.1 m/s. Reaction time was not computed for a small minority of trials (1260 out of 41560) where the velocity did not exceed 0.1 m/s. Peak velocity was defined as the cursor’s highest tangential velocity. We computed the tangential velocity by linearly resampling the cursor’s position at the times recorded by the Flock of Birds and computing the distance traversed by the cursor between two consecutive samples divided by the time elapsed. Resampling was necessary because, occasionally, the recorded time at which a sample was collected by the Flock of Birds did not match the true time at which it was collected, causing the calculated velocity to be inaccurate. Velocity profiles were also smoothed using a third-order Savitzky-Golay filter.

Initial reach direction was defined as the direction of the instantaneous velocity vector 150 ms after movement initiation. Initial reach direction error was computed as the difference in angle between this instantaneous velocity vector and the vector pointing from the target on the previous trial to the target on the current trial. Probability density functions were estimated for reach direction errors using a kernel smoothing function, implemented as the ksdensity function in MATLAB. We measured the variability in participants’ initial reach direction errors (i.e., how consistently straight participants reached towards the target) by fitting a mixture model to this data. In the model, we assumed that participants’ reach direction errors, *x*, were generated by one of two causes: 1) an error from a goal-directed reach towards the target (modeled as a von Mises distribution) or 2) an error from a reach in a random direction (modeled as a uniform distribution). The probability density function of the mixture model, mix(·), was defined as

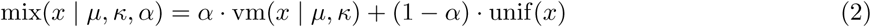

where *α* is a parameter valued between 0 and 1 weighting the probability density functions of the von Mises, vm(·), and uniform distributions, unif(·). The probability density functions of the individual distributions were defined as

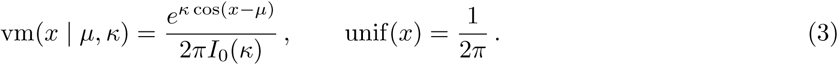

Here, *µ* and *κ* are the mean and concentration of the von Mises distribution and *I*_0_(*·*) is the modified Bessel function of the first kind with order 0.

The parameters *µ, κ*, and *α* were fit to the data from single participants in each block via maximum likelihood estimation. Specifically, we used the MATLAB function fmincon to determine the values of the parameters that would maximize the below likelihood function over the *n* trials within one block:

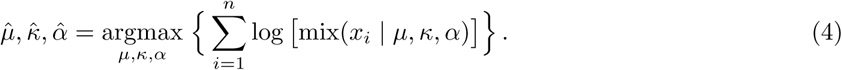

Then, using the fitted concentration parameter of the von Mises distribution, 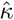, we computed the circular standard deviation, *σ*, as

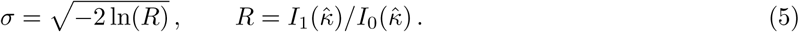

We used *σ* as our measure of the variability of participants’ reach direction errors.

To assess the whether participants exhibited habitual behavior during the flip block, we quantified each participant’s tendency to reach towards the true target versus a virtual target flipped across the mirroring axis. More specifically, we assumed that for each trial, participants’ initial reach direction could be explained by at least one of three causes: 1) a goal-directed reach towards the target, 2) a habitual reach towards the mirrored target, and 3) a reach aimed towards neither target (i.e., random movement). We modeled the first two causes as von Mises distributions with different means—*ϕ*_*a*_ and *ϕ*_*m*_, set by the direction of the actual and mirrored targets, respectively—but the same concentration parameter, *κ*. For each participant, we fixed the concentration parameter to be equal to the 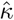’s estimated for late learning in Eq 4. We modeled the third component, random movements, as a uniform distribution.

Assuming that each participant’s behavior within one block could be modeled as a weighted mixture of these three distributions, mix^*′*^(·), we used the MATLAB function fmincon to determine the weights, *α*_*a*_ and *α*_*m*_, that would maximize the following likelihood function over the *n* trials within one block:

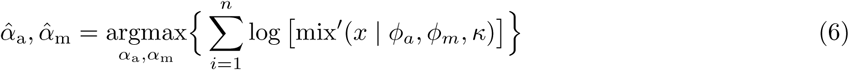

Where

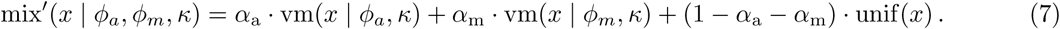

Here, *x* represents participants’ reach directions while *α*_a_ and *α*_m_ correspond to the probabilities that a participant reached towards the actual, and mirrored targets, respectively. Definitions for vm(·) and unif(·) can be found in Eq 3. We used 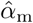 as our metric for the strength of habitual behavior. For Fig 4E, instead of fitting this model to all trials in the flip block, we fit the model to either the first or second half of trials in this block.

We used the fitted weights from this approach to classify each trial as either goal-directed, habitual, or random. For each trial, we computed the probability that the reach direction was generated from each of the three mixture components under the fitted mixture model’s probability density function (in essence, computing *p*(reach direction | goal-directed), *p*(reach direction | habitual), and *p*(reach direction | random). Trials were classified as goal-directed, habitual, or random based on which of these three probabilities was the highest. We excluded trials where the direction of the target was within 30^*°*^ of the mirroring axis (the *y*-axis) as the von Mises distributions for the goal-directed versus habitual reaches would have similar means and, therefore, would be too difficult to distinguish from each other (1153 out of 3200 data points excluded). We used this classification to compute the reaction times of goal-directed versus habitual reaches in Fig 4D.

Additionally, we compared the model in Eq 7 with an alternative model which was designed to capture behavior where participants would move their right hand (controls vertical cursor movement) in the correct direction but their left hand (controls horizontal cursor movement) would generate random movements. We modeled participants reach directions, *x*, given the target’s direction, *ϕ*_*a*_, as a mixture of two weighted uniform distributions:

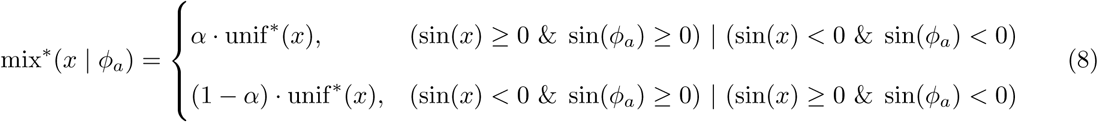

Where

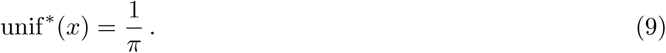

Here, *α* is the probability that the cursor moved vertically in the correct direction. The fits for the models in Eq 7 and Eq 8 were compared using BIC. Model recovery analyses were performed by simulating data from both of these models, fitting both models to each simulated dataset, comparing fits using BIC, and generating a confusion matrix. To generate data from Eq 7, we used values for *α*_*a*_ and *α*_*m*_ that ranged between 0 and 1, and we fixed *κ* = 3. Lower *κ*’s set higher variability for the von Mises distributions (i.e., harder to distinguish from the model in Eq 8), so we fixed *κ* to be the lowest average *κ* that we observed in the late learning data from any group, as estimated in Eq 4. Model recovery across different choices of parameters were compared by computing the accuracy of the confusion matrices.

As an alternative approach to assessing whether participants exhibited habitual behavior while using the flipped mapping, we assessed whether their horizontal cursor movements were aimed away from the target. Using only the cursor’s *x*-axis position, for each trial, we determined whether the cursor’s instantaneous velocity vector was aimed towards the right or left 150 ms after the cursor deviated 1 cm from the center of the starting target (i.e., the radius of the target). We classified cursor movements in each trial as moving away from the target if the velocity vector’s direction was opposite of the direction of the target relative to the starting position (e.g., target located to the left but cursor moving towards the right). This method was unable to compute an initial horizontal reach direction on a small minority of trials (95 out of 3200 trials) where the target on the current trial was either directly above or below the target from the previous trial. This was because either: 1) the cursor did not deviate 1 cm horizontally away from the center of the starting target (i.e., the radius of the target), making it impossible to detect the time of movement initiation, or 2) the detected movement initiation time was less than 150 ms prior to the end of the trial, meaning that the trial ended before the time at which we assessed reach direction. These trials were excluded from the analysis.

### Analysis of tracking task

Data from two tracking trials (each from different subjects) were excluded from the analysis because our experiment hardware failed to accurately record the positions of stimuli with known positions. Tracking error was computed as the mean-squared error between the cursor’s and target’s positions. Time-domain trajectories of the cursor’s and target’s position (the first 60 seconds of each trial following the initial 5 second ramp period) were converted to phasors (complex numbers representing sinusoids) in the frequency domain via the discrete Fourier transform. An input-output transfer function was computed at every frequency by dividing the cursor’s phasor by the target’s phasor. This transfer function described the relationship between the cursor and target sinusoids in terms of gain (relative amplitude) and phase (difference in time).

Using these transfer functions, we sought to describe the direction that participants moved their cursor to track the target. In this task, participants’ cursor movements would conventionally be described as phase lagged relative to the target with a positive gain (i.e., moving in the same direction as the target with a time delay). However, when a mirror reversal has been applied (such as in the flipped mapping), participants’ may habitually continue to use their original control policy, causing their movements to be flipped across the mirroring axis relative to before. Although the relationship between movements before and after the flip could be described as movements with positive gain but now in antiphase (i.e., moving in the same direction as the target but with more time delay), a better way to describe them would be to say that the movements have the same phase but a negative gain (i.e., moving with the same time delay but in the opposite direction of the target).

Given that conventional analysis methods always yield a positive gain to describe frequency-domain data, we used the method described in [37] to compute a signed gain, *g*, relating cursor and target movements. This was computed as the dot product between transfer functions:

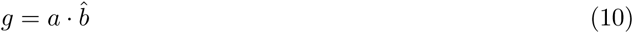

where *a* is the transfer function for a given block of interest and 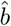 is the transfer function at baseline with unit length. Computing the dot product implicitly fixes the phase of cursor movements to be the same as baseline across all blocks, allowing a signed gain to be computed. This assumption of fixed phase is valid for analyzing data in late learning as participants’ phase lags under the bimanual mapping became more similar to baseline through practice. We computed this signed gain between each axis of target and cursor movement (*x*-axis target movement and *x*-axis cursor movement, *x*-axis target movement and *y*-axis cursor movement, etc.), building a series 2 *×* 2 matrices relating the transformation between the two trajectories where each matrix represented the transformation within a small bandwidth of frequencies. The green (purple) arrows in the Fig 3C and 5A were generated by plotting the first (second) column of each matrix. Fig 3D and Supplementary Fig 5B was generated by plotting the element in the first row and first column of each matrix.

To quantify the strength of habitual behavior in Fig 5B, we reanalyzed the gain between *x*-axis target and *x*-axis cursor movements from the flip blocks by fixing their phases to be the same as late learning. We did this because any habitual behavior would manifest as lingering usage of the originally learned bimanual mapping, and the habit should be measured with respect to behavior under this mapping. When analyzing these normalized gains at the group level, we excluded one outlier participant in the 10-day group who exhibited dramatically more negative gains than other participants within the group (lower left participant in Supplementary Fig 4). To compare the habitual behavior we observed between the point-to-point and tracking tasks, we correlated each participant’s *α*_*m*_ from Eq 6 with their normalized gains (averaged over the highest three frequencies) from Fig 5B via linear regression.

### Statistics

Most primary statistical analysis were performed by fitting linear mixed effects models to the data. For all analyses in Fig 2, the models used group (2-, 5-, or 10-day) and block (2-day: day 1 vs. day 2; 5-day: day 2 vs. day 5; 10-day: day 5 vs. day 10) as fixed effects and subject as a random effect. For Fig 4C and E, models used the same group and subject effects but with a different set of blocks being compared ([late learning vs. flip block] and [first half of flip block vs. second half of flip block], respectively). For Fig 4D, models used the same group and subject effects but with reach type as an additional fixed effect (goal-directed vs. habitual). Post-hoc pairwise comparisons were performed using the Tukey test.

For data from the tracking task, mixed effects models were fit using the same effects as Fig 2 but with an additional fixed effect of frequency. We also fit separate models to data from each frequency because behavior varied dramatically as a function of frequency. Post-hoc pairwise comparisons were performed using the Tukey test. An additional Bonferroni correction factor of 6 was applied to the p-values for pairwise comparisons to account for the separate models fits for each frequency. Additionally, to determine whether participants exhibited significantly negative gains in the tracking task, for each frequency, we performed a series of one-sample t-tests and corrected for multiple (6) comparisons using a Holm-Bonferroni correction with *α* = 0.05.

### Data and Code Availability

The data and code used in the study can be found in the Johns Hopkins University Data Archive: https://doi.org/10.7281/T1/FWDYPW.

## Acknowledgements

We thank Yue Du, John Krakauer, Amy Bastian, and Christopher Fetsch for helpful discussions. This material is based upon work supported by the National Science Foundation under Grant No. 1825489 (N.J.C.). This work was also supported by Facebook Technologies (A.M.H. and C.S.Y.). C.S.Y was supported by the Link Foundation Modeling, Simulation & Training Fellowship.

## Author Contributions

C.S.Y and A.M.H. conceptualized the experiment. N.J.C provided technical expertise for the tracking methodology. C.S.Y collected data and performed data analysis while A.M.H. and N.J.C. supervised. C.S.Y. and wrote the manuscript while all authors reviewed and edited the manuscript.

## Supplementary Material

**[Supplementary Fig 1].**
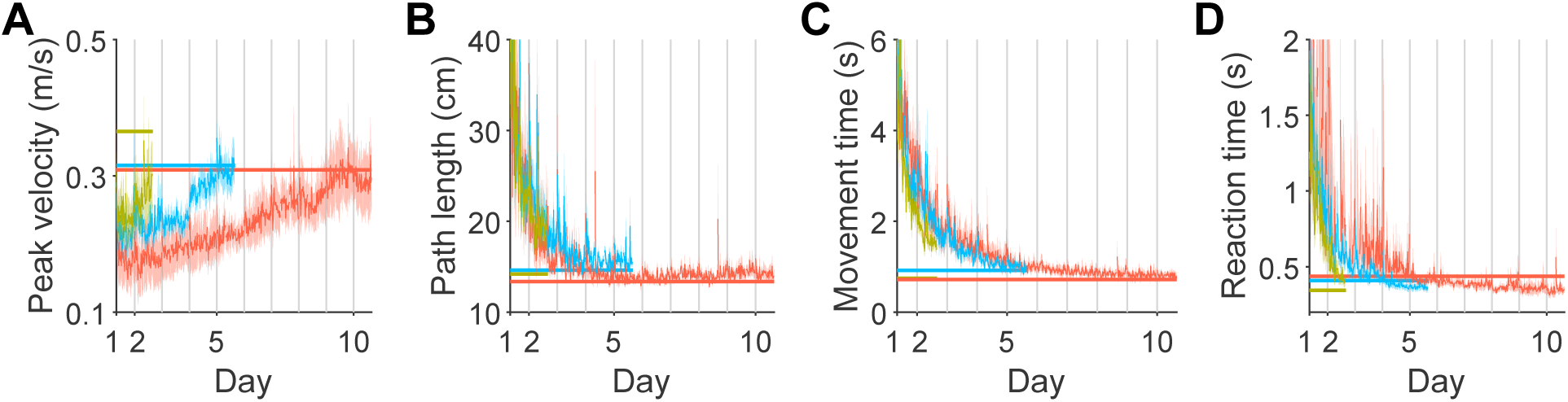
Kinematics of point-to-point movements. Peak velocity, path length, movement time, and reaction time of point-to-point movements from baseline to late learning. Data were averaged across bins of 5 trials and error bars indicate SEM across participants. Baseline values for each group are shown as horizontal lines. Different days are demarcated by gray vertical lines.

**[Supplementary Fig 2].**
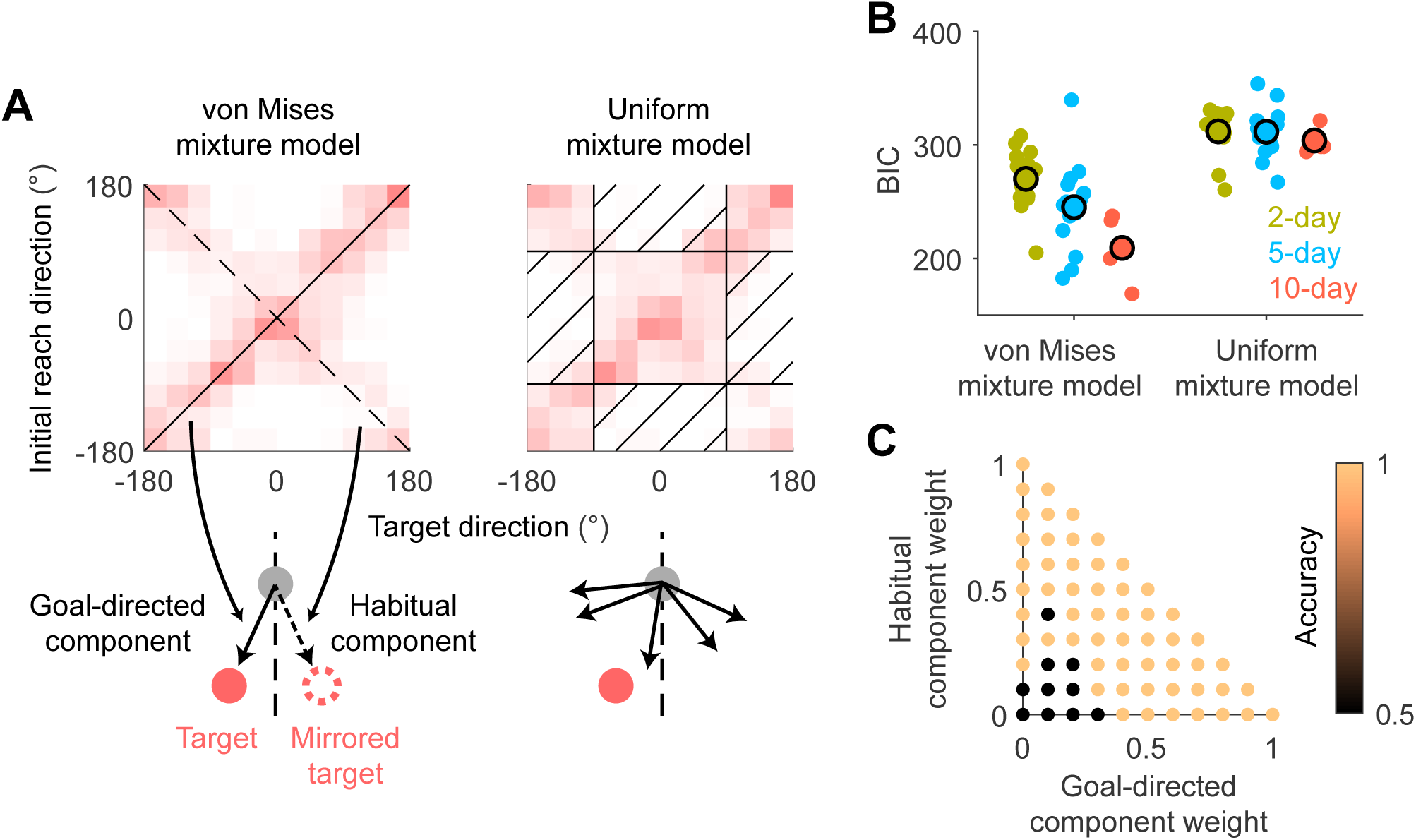
Model comparison and model recovery analysis. A: Depictions of the two mixture models fit to participants’ initial reach directions. (Left) Model composed of two von Mises distributions centered around the *y* = *x* (captures goal-directed reaches; solid line) and *y* = −*x* lines (captures habitual reaches; dashed line) as well as a uniform distribution (captures random reaches). This model would be appropriate if cursor movements were generally aimed either directly towards the target or towards a virtual target mirrored across the *y*-axis. (Right) Model composed of two weighted uniform distributions capturing cursor movements towards (non-hatched area) or away (hatched area) from the target in the *y*-axis. This model would be appropriate if participants generated correct right-hand movements (vertical movements of the cursor) but random left-hand movements (horizontal movements of the cursor). Heat maps of data from the 2-day group in the flip block are shown for reference. 0^*°*^ is defined as the positive *y*-axis. B: Comparison of models fitted to data from the flip block. Lower BIC scores indicate better fits. C: Model recovery analysis. We simulated data from both the von Mises and uniform mixture models and then fit both models to this data, examining which model fit the data better using a confusion matrix. Accuracy of the confusion matrix is plotted for these combinations of weights. When data are generated from the von Mises mixture model with the weights of the two von Mises components summing to be over 0.5, the accuracy was 100%. Empirically, we found that the sum of these weights is generally well over 0.5 (Fig 4D), suggesting that the von Mises mixture model can be recovered with the type of behavior we observed.

**[Supplementary Fig 3].**
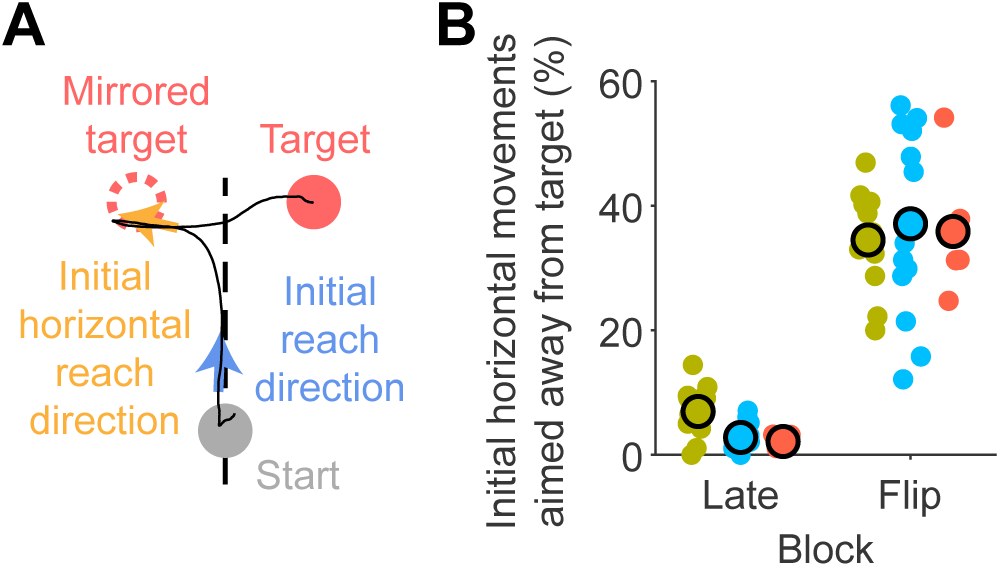
Reach direction analysis of horizontal cursor movements. The analysis in Fig 4C quantified habitual behavior based on the cursor’s *initial* reach direction, which may not have captured habitual behavior which occurred *later* in the trial. As a result, we performed an additional analysis quantifying the proportion of horizontal cursor movements initially aimed away from the target. A: Example cursor trajectory (black line) where habitual behavior occurred towards the end of a reach; although the cursor’s movement was initially aimed along the mirroring axis (blue), the cursor was aimed away from the true target towards the end of the movement (orange). Mirroring axis shown as a dashed line. B: Percentage of trials where the cursor’s initial horizontal movement was aimed away from the target.

**[Supplementary Fig 4].**
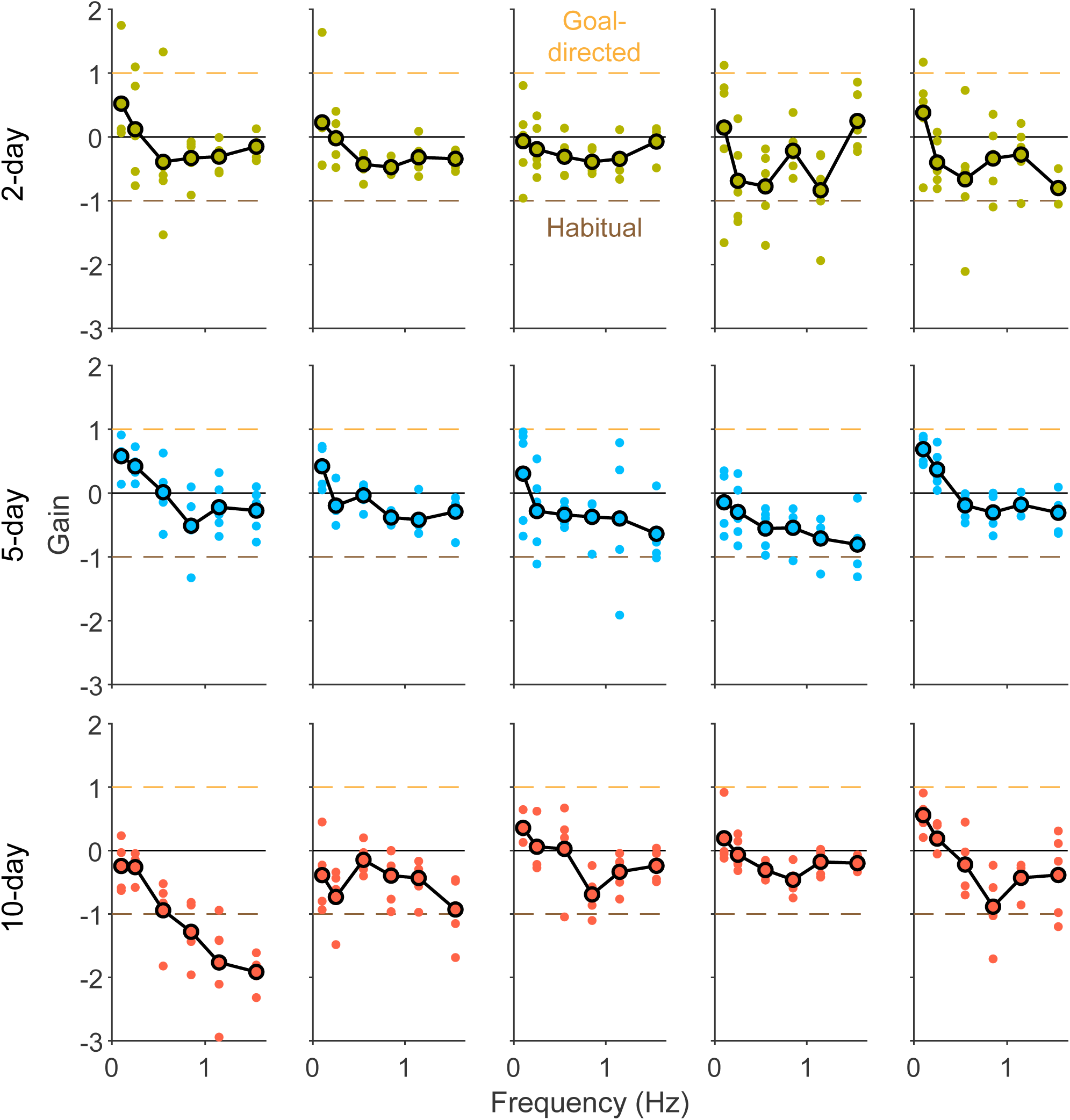
Gain of movements under the flipped mapping from single participants. Data from five participants in each group during the first flipped tracking block are shown, with individual trial data shown as small dots and averages across trials shown as black lines. These data were used to generate Fig 5B.

**[Supplementary Fig 5].**
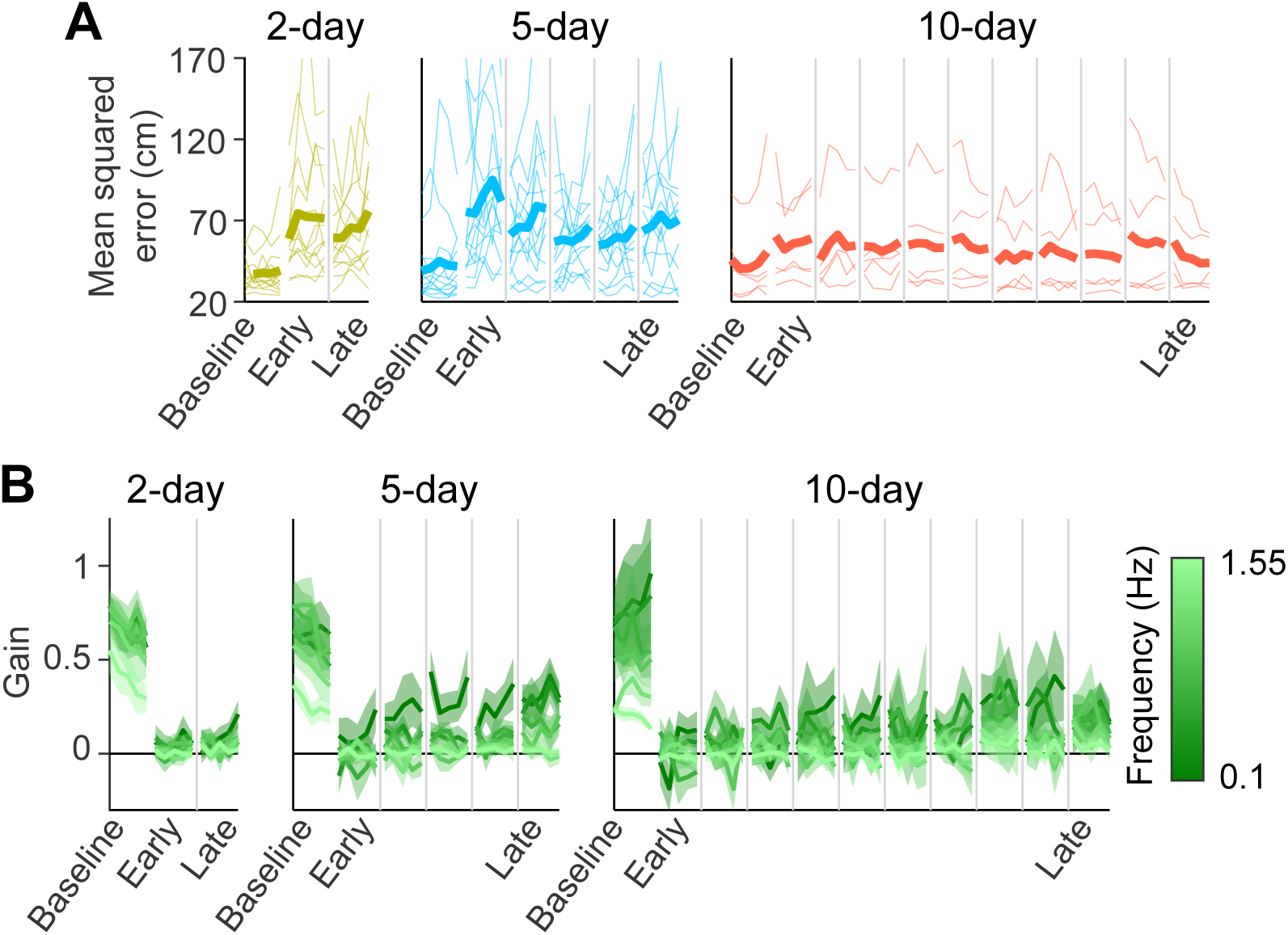
Analysis of tracking trials without visual feedback of the cursor. A: Mean-squared error between cursor and target positions. Thin lines indicate individual participants while thick lines indicate group averages. Data collected from different days are separated by gray lines. B: Gain between target and cursor movements at frequencies of *x*-axis target movement. Lower and higher frequencies are depicted as darker and lighter colors, respectively. Error bars indicate SEM across participants.

## References

[1] Sandlin, D. The backwards brain bicycle - smarter every day 133 (2015). URL https://www.youtube.com/watch?v=MFzDaBzBlL0&t=2s.

[2] Wood, W. & Neal, D. T. A new look at habits and the habit-goal interface. Psychol. Rev. 114, 843–863 (2007).

[3] Seger, C. & Spiering, B. A critical review of habit learning and the basal ganglia. Front. Syst. Neurosci. 5 (2011).

[4] Hélie, S. & Cousineau, D. The cognitive neuroscience of automaticity: Behavioral and brain signatures, 141–159 (Nova Science Publishers, 2014).

[5] Adams, C. D. & Dickinson, A. Instrumental responding following reinforcer devaluation. Q. J. Exp. Psychol. Sect. B 33, 109–121 (1981).

[6] Balleine, B. W. & O’Doherty, J.P. Human and rodent homologies in action control: Corticostriatal determinants of goal-directed and habitual action. Neuropsychopharmacology 35, 48–69 (2010).

[7] de Wit, S. et al. Shifting the balance between goals and habits: Five failures in experimental habit induction. J. Exp. Psychol. 147, 1043–1065 (2018).

[8] Luque, D., Molinero, S., Watson, P., López, J., Francisco & Le Pelley, M. E. Measuring habit formation through goal-directed response switching. J. Exp. Psychol. Gen. 149, 1449–1459 (2020).

[9] Ceceli, A. O., Myers, C. E. & Tricomi, E. Demonstrating and disrupting well-learned habits. PLoS One 15, 1–28 (2020).

[10] Popp, N. J., Yokoi, A., Gribble, P. L. & Diedrichsen, J. The effect of instruction on motor skill learning. J. Neurophysiol. 124, 1449–1457 (2020).

[11] Tricomi, E., Balleine, B. W. & O’Doherty, J.P. A specific role for posterior dorsolateral striatum in human habit learning. Eur. J. Neurosci. 29, 2225–2232 (2009).

[12] McDonald, R. J., King, A. L. & Hong, N. S. Context-specific interference on reversal learning of a stimulus-response habit. Behav. Brain Res. 121, 149–165 (2001).

[13] Faure, A., Haberland, U., Condé, F. & Massioui, N. E. Lesion to the nigrostriatal dopamine system disrupts stimulus-response habit formation. J. Neurosci. 25, 2771–2780 (2005).

[14] Yin, H. H. & Knowlton, B. J. The role of the basal ganglia in habit formation. Nat. Rev. Neurosci. 7, 464–476 (2006).

[15] Robbins, T. W. & Costa, R. M. Habits. Curr. Biol. 27, R1200–R1206 (2017).

[16] Salmon, D. P. & Butters, N. Neurobiology of skill and habit learning. Curr. Opin. Neurobiol. 5, 184–190 (1995).

[17] Bernacer, J. & Murillo, J. I. The aristotelian conception of habit and its contribution to human neuroscience. Front. Hum. Neurosci. 8 (2014).

[18] Graybiel, A. M. & Grafton, S. T. The striatum: Where skills and habits meet. Cold Spring Harb. Perspect. Biol. 7, a021691 (2015).

[19] Haith, A. M. & Krakauer, J. W. The multiple effects of practice: skill, habit and reduced cognitive load. Curr. Opin. Behav. Sci. 20, 196–201 (2018).

[20] Marien, H., Custers, R. & Aarts, H. Understanding the Formation of Human Habits: An Analysis of Mechanisms of Habitual Behaviour, 51–69 (Springer International Publishing, Cham, 2018). URL https://doi.org/10.1007/978-3-319-97529-0_4.

[21] Du, Y., Krakauer, J. W. & Haith, A. M. The relationship between skills and habits in humans. PsyArXiv (2021).

[22] Gritsenko, V. & Kalaska, J. F. Rapid online correction is selectively suppressed during movement with a visuomotor transformation. J. Neurophysiol. 104, 3084–3104 (2010).

[23] Telgen, S., Parvin, D. & Diedrichsen, J. Mirror reversal and visual rotation are learned and consolidated via separate mechanisms: Recalibrating or learning de novo? J. Neurosci. 34, 13768–13779 (2014).

[24] Collins, D., Morriss, C. & Trower, J. Getting it back: A case study of skill recovery in an elite athlete. Sport Psychol. 13, 288–298 (1999).

[25] Hanin, Y., Malvela, M. & Hanina, M. Rapid correction of start technique in an olympic-level swimmer: A case study using old way/new way. J. Swim. Res. 16, 11–17 (2004).

[26] Carson, H. J., Collins, D. & Jones, B. A case study of technical change and rehabilitation: Intervention design and multidisciplinary team interaction. Int. J. Sport Psychol. 45, 57–78 (2014).

[27] Haith, A. M., Yang, C. S., Pakpoor, J. & Kita, K. De novo motor learning of a bimanual control task over multiple days of practice. bioRxiv (2021).

[28] Krakauer, J. W., Pine, Z. M., Ghilardi, M.-F. & Ghez, C. Learning of visuomotor transformations for vectorial planning of reaching trajectories. J. Neurosci. 20, 8916–8924 (2000).

[29] Fernandez-Ruiz, J., Wong, W., Armstrong, I. T. & Flanagan, J. R. Relation between reaction time and reach errors during visuomotor adaptation. Behav. Brain Res. 219, 8–14 (2011).

[30] Bond, K. M. & Taylor, J. A. Flexible explicit but rigid implicit learning in a visuomotor adaptation task. J. Neurophysiol. 113, 3836–3849 (2015).

[31] Morehead, J. R., Qasim, S. E., Crossley, M. J. & Ivry, R. Savings upon re-aiming in visuomotor adaptation. J. Neurosci. 35, 14386–14396 (2015).

[32] Hardwick, R. M., Forrence, A. D., Krakauer, J. W. & Haith, A. M. Time-dependent competition between goal-directed and habitual response preparation. Nat. Hum. Behav. 3, 1252âĂŞ–1262 (2019).

[33] Roth, E., Zhuang, K., Stamper, S. A., Fortune, E. S. & Cowan, N. J. Stimulus predictability mediates a switch in locomotor smooth pursuit performance for eigenmannia virescens. J. Exp. Biol. 214, 1170–1180 (2011).

[34] Madhav, M. S., Stamper, S. A., Fortune, E. S. & Cowan, N. J. Closed-loop stabilization of the Jamming Avoidance Response reveals its locally unstable and globally nonlinear dynamics. J. Exp. Biol. 216, 4272–4284 (2013).

[35] Sponberg, S., Dyhr, J. P., Hall, R. W. & Daniel, T. L. Luminance-dependent visual processing enables moth flight in low light. Science 348, 1245–1248 (2015).

[36] Zimmet, A. M., Cao, D., Bastian, A. J. & Cowan, N. J. Cerebellar patients have intact feedback control that can be leveraged to improve reaching. eLife 9, e53246 (2020).

[37] Yang, C. S., Cowan, N. J. & Haith, A. M. De novo learning versus adaptation of continuous control in a manual tracking task. eLife 10, e62578 (2021).

[38] Daw, N. D., Niv, Y. & Dayan, P. Uncertainty-based competition between prefrontal and dorsolateral striatal systems for behavioral control. Nat. Neurosci. 8, 1704–1711 (2005).

[39] Dolan, R. J. & Dayan, P. Goals and habits in the brain. Neuron 80, 312–325 (2013).

[40] Hogarth, L. A Critical Review of Habit Theory of Drug Dependence, 325–341 (Springer International Publishing, Cham, 2018). URL https://doi.org/10.1007/978-3-319-97529-0_18.

[41] Maisto, D., Friston, K. & Pezzulo, G. Caching mechanisms for habit formation in active inference. Neurocomputing 359, 298–314 (2019).

[42] Gläscher, J., Daw, N., Dayan, P. & O’Doherty, J. P. States versus rewards: Dissociable neural prediction error signals underlying model-based and model-free reinforcement learning. Neuron 66, 585–595 (2010).

[43] Toner, J., Montero, B. G. & Moran, A. The perils of automaticity. Rev. Gen. Psychol. 19, 431–442 (2015).

[44] R Core Team. R: A Language and Environment for Statistical Computing. R Foundation for Statistical Computing, Vienna, Austria (2020). URL https://www.R-project.org/.

[45] Bates, D. & Maechler, M. Matrix: Sparse and Dense Matrix Classes and Methods (2019). URL https://CRAN.R-project.org/package=Matrix. R package version 1.2-18.

[46] Bates, D., Mächler, M., Bolker, B. & Walker, S. Fitting linear mixed-effects models using lme4. J. Stat. Softw. 67, 1–48 (2015).

[47] Kuznetsova, A., Brockhoff, P. B. & Christensen, R. H. B. lmerTest package: Tests in linear mixed effects models. J. Stat. Softw. 82, 1–26 (2017).

[48] Lenth, R. emmeans: Estimated Marginal Means, aka Least-Squares Means (2020). URL https://CRAN.R-project.org/package=emmeans. R package version 1.4.8.

